# Lag Effects in Primate Brain Size Evolution: A Re-Evaluation

**DOI:** 10.1101/355453

**Authors:** R.I.M. Dunbar

## Abstract

The question as to whether there is a lag between brain and body mass evolution was ostensibly solved two decades ago by Deaner & Nunn (1999) who used phylogenetic methods to show that there was no evidence to suggest that changes in brain size lagged behind changes in body size. However, their assumption that body size would always change ahead of brain size is open to question. In addition, many of their datapoints are confounded by grade shift effects. A reanalysis of their data controlling for these confounds shows that there is in fact a strong lag effect, but that the direction of the lag is the reverse of that originally assumed: brain size typically changes first, and does so under selection from changes in group size. The data suggest that it takes about 2.0 million years for body size to converge back onto the conventional allometric relationship with brain size. In the meantime, species that have increased brain size are likely to incur a significant energy cost that must be met from elsewhere. I show that they seem to do so by changing to a more nutrient-rich diet.

## 1. Introduction

Ever since Jerison’s (1973) seminal analyses, it has been known that, across mammals in general and primates in particular, brain size is correlated with body mass in an allometric power relationship. Nevertheless, there is considerable variation around the common regression line, and it was commonly suggested that this is a consequence of a lag effect in which brain mass takes time to catch up with changes in body mass (Jerison 1973; Lande 1979; Martin & Harvey 1985; Willner & Martin 1985; Deacon 1990b, Deacon 1997; Aboitiz 1996). This assumption is based mainly on the fact that body size is relatively labile, and can vary considerably within species as a function of local environmental conditions (Dunbar 1990; Bettridge et al. 2010), whereas the complex interconnectivity of brain systems means that it is likely to take longer to bring together the necessary genetic changes without disrupting functional neural systems. Such an effect might explain the well established fact that body size has outstripped brain size in most domesticated species (Hemmer 1990). It is, howevr, assumed that, given enough time, brain and body size converge on the common regression line under pressure from some form of stabilising selection. That brain and body mass are not yoked in close genetic linkage is confirmed by breeding experiments showing that brain size and body size can undergo independent selection, at least in the short term (Riska & Atchley 1985).

Deaner & Nunn (1999) developed a novel method for testing the lag hypothesis that involved plotting the residuals of phylogenetic contrasts in brain mass regressed on contrasts in body mass for a sample of primates against date of divergence. They tested the explicit hypothesis that the lag would be directional: body mass would always change first (hence this was always taken as a positive change), and the lag would thus necessarily be brain mass lagging behind body mass. This being so, a lag, if present, should be evidenced by a positive correlation because ‘young’ nodes would consist of species pairs with negative residuals when body mass contrasts are constrained always to be positive. They found that there was no correlation between the two variables, either for males or for females, or when controlling for ecology, and concluded that there was no evidence for lag effect.

However, there is no principled reason why body size has to change first. Much will depend on the ecological pressures acting on brain and body mass, and these can be very different. Brain size is known to be driven by changes in the cognitive demands imposed by increasing social group size (Dunbar 1998; Perez-Barberia et al. 2007; Shultz & Dunbar 2007; Dunbar & Shultz 2007, 2010). If brain size is responding mainly to pressures to evolve larger group sizes but body size responds mainly to ecological pressures (e.g. nutrient availability), then the two need not be in close linkage. A further problem that emerges with their analysis is that many of the nodes they use are not closely related and involve major grade shifts, potentially resulting in further confounds. In this paper, I reanalyse their data and show that there is, in fact, clear evidence for a lag effect.

## 2. Methods

In order to ensure that any differences between Deaner & Nunn’s original results and the new analyses are due to methodology and not to different data samples, I use the dataset provided by Deaner & Nunn (1999). Their data are CAIC contrasts without reference to divergence times on the grounds that they wanted to use divergence time as an independent variable in the analysis. They explicitly used only nodes that were tip comparisons (i.e. comparisons between living species) and avoided nodes at higher levels in the phylogeny (which hence have to be estimated). They used the Stephan et al. (1969) brain dataset because this provides data on actual brain and body masses for the same individual specimens. This dataset yields a set of 24 nodes for which contrasts in brain mass and contrasts in body mass are available. Although a much larger sample of species is available for cranial volumes (e.g. Isler et al. 2008), Deaner & Nunn (1999) argued (rightly) that these datasets risked introducing unnecessary error variance: their main problem is that the body mass and cranial volume data derive from different animals. I did, however, check out this data source, but the sample size is no larger once the data are filtered for group size and divergence dates. I use the divergence times as given by Deaner & Nunn (1999) (based on Purvis 1995) as well as more recent estimates provided by Perelman et al. (2011). Group size and dietary data for individual species are from Campbell et al. (2007), with the exception of baboon group sizes which derive from Bettridge et al. (2010).

Where we are testing a directional hypothesis, one-tailed statistical tests are appropriate: in such cases, a significant correlation in the opposite direction would be evidence *against* the hypothesis. In testing all other hypotheses, 2-tailed tests are used.

## 3. Results

Fig. 1 plots the residuals in the contrasts for brain mass as a function of body mass against divergence time. Essentially these are the data as presented by Deaner & Nunn (1999) and are the data they use to justify their conclusion. Deaner & Nunn (1999) tested for a positive relationship with time, but found a regression slope that did not differ significantly from *b*=0, and concluded from this that there was no evidence for any lag effect. However, inspection of Fig. 1 indicates that the datapoints are evenly distributed either side of zero, with a variance that has a funnel shape, so it is no surprise that there was the correlation did not differ from zero: this is simply a consequence of using residuals from the common regression. More importantly, a funnel-shaped pattern of variance is exactly what would be expected if there is a lag effect but the direction of the lag is unspecified (i.e. sometimes body size changes first, and sometimes brain size changes first).

**Fig. 1.**
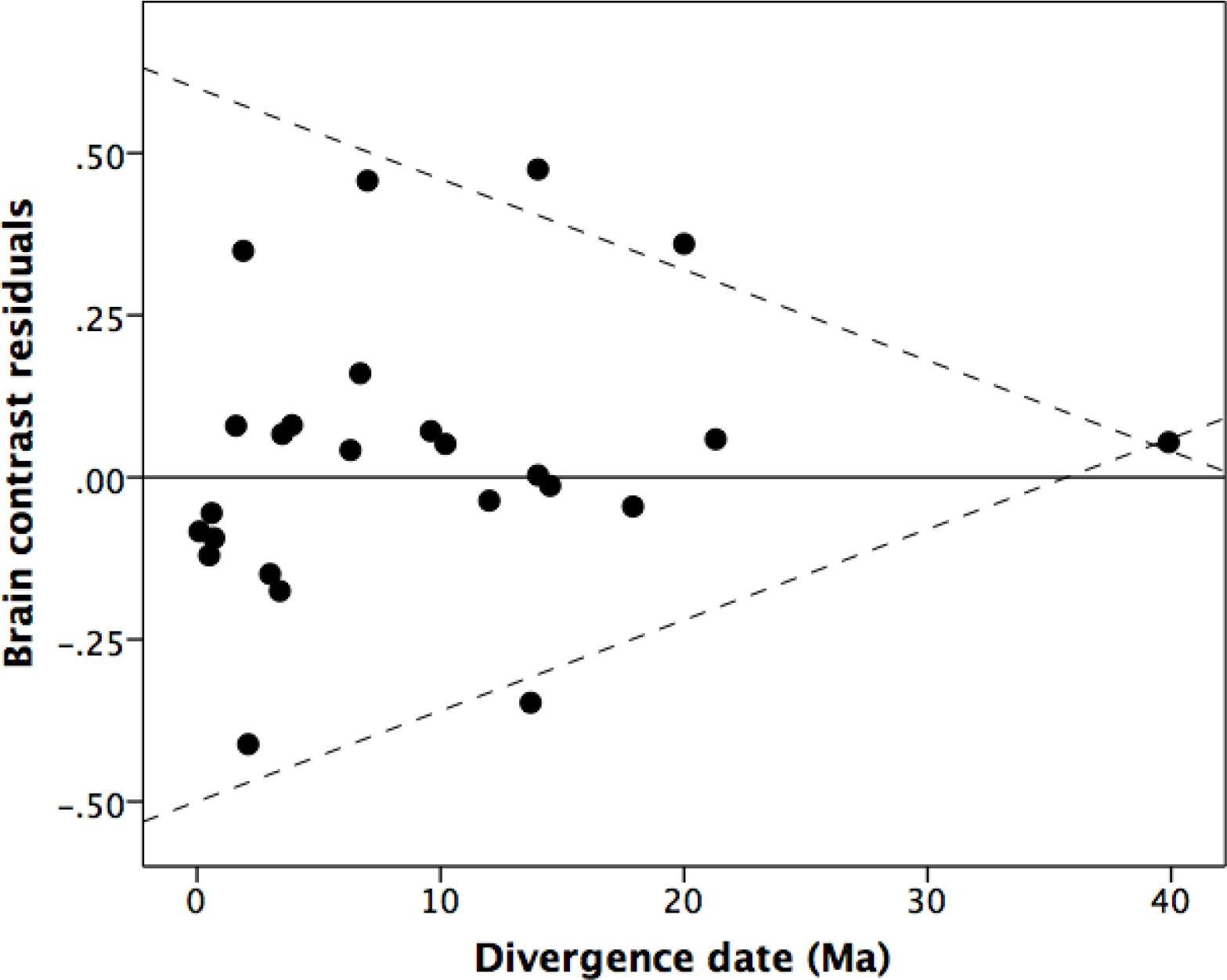
Residuals of contrast in brain mass regressed against contrasts in brain mass plotted against time of divergence for the species pairs. Dashed lines are the upper and lower bounds.

This being so, absolute residuals should be plotted against divergence date (Fig. 2). The relationship is clearly negative, albeit not significant (Spearman r_s_=-0. 074, p=0.362 1-tailed). One obvious problem with these data is that they mix contrasts of very different taxonomic status, namely contrasts between closely related species (i.e. those within the same genus) and species that belong to different genera, and in some cases even different families (highlighted as solid and open symbols). Most of the latter are strepsirrhines, but they also include the contrast between *Homo* and *Pan* and that between *Allouatta* and *Lagothrix*, both of which involve very large differences in sociality (especially group size) as well as brain volume. These risk confounding the lag relationship with major grade shifts in brain size (see Aiello & Dunbar 1992; Dunbar 1993). This is especially problematic for the strepsirrhines, where there are grade shifts between nocturnal and diurnal species (Barton 1998), as well as the grade shift between strepsirrhines and haplorhines. They also all entail large differences in sociality involving the transition between semi-solitary species and those that live in multimale/multifemale groups. If we consider only pairwise comparisons for closely related species (i.e. those belonging to the same genus), the correlation between residual brain mass and divergence date is in fact highly significant (*r*_*s*_=-0.593, N=13, p=0.017 1-tailed). In contrast, the correlation for the between-genus contrasts is not significant (Spearman *r*=-0.210, N=12, p=0.256 1tailed), although the slope is similar to that for within-genus nodes.

**Fig. 2.**
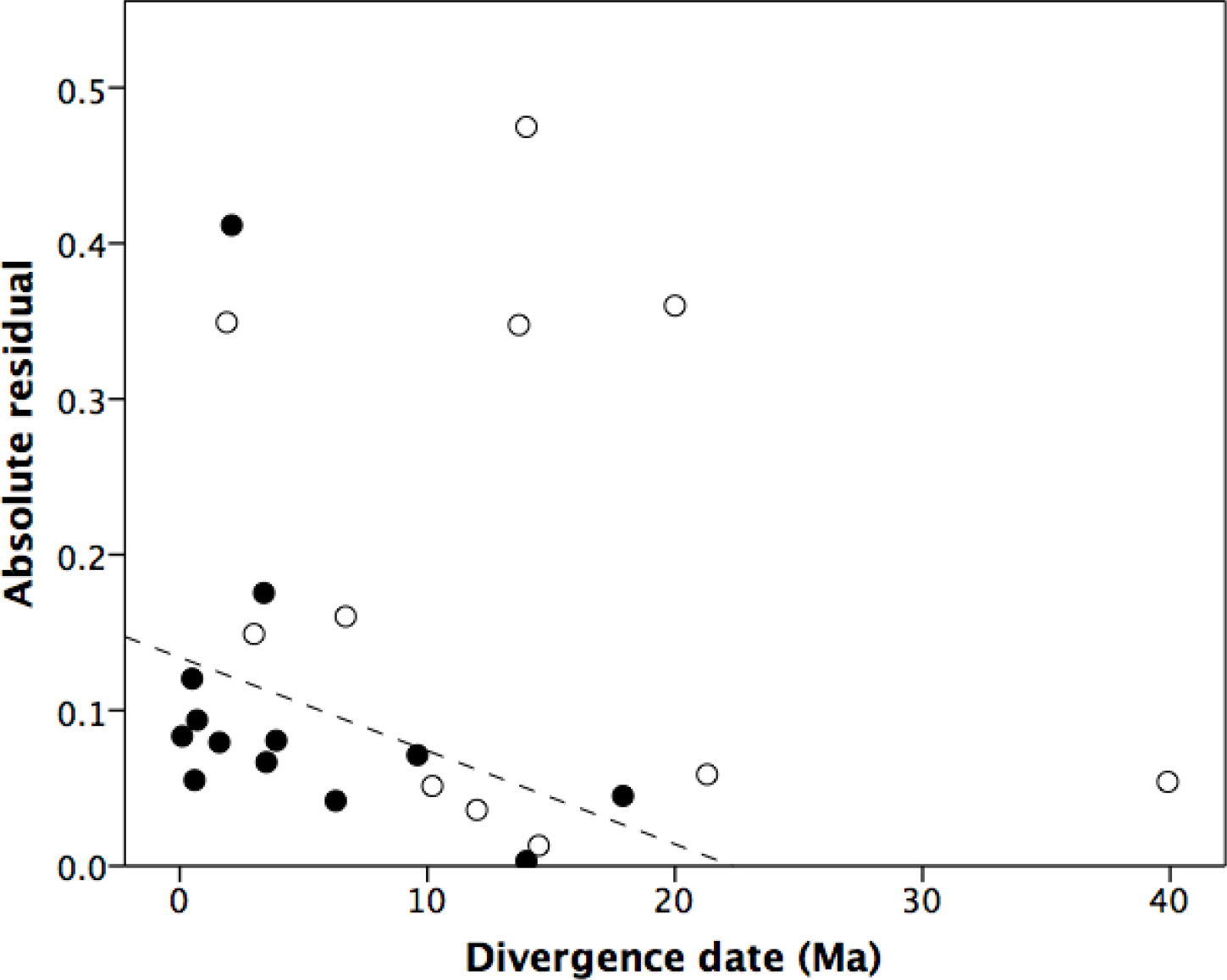
Absolute residuals of contrast in brain mass, plotted against time since divergence. Dashed line is the least squares regression for these data points. Data points from Fig. 1.

Deaner & Nunn (1999) tested for a relationship between social group size and residual brain size, but found none. However, using more up-to-date data on species mean group sizes, there is in fact a significant linear regression between contrasts in mean taxon group size and residuals in brain size contrasts (Fig. 3; F_1,17_=6.15, r^2^=0.266, p=0.024 2-tailed). Two things are immediately apparent, however: first, there is a very striking grade difference between strepsirrhines (prosimians) and haplorhines (anthropoids) and, second, the relationship is clearly non-linear (hence, the likely reason why Deaner & Nunn obtained a non-significant result when using a linear regression). Both effects are in fact well established features of the social brain hypothesis (Dunbar 1993, 1998), and were well known at the time. Partitioning the data by sub-order, and logging group size, yields significant improvements in fit (prosimians: F_1,2_=7.96, r^2^=0.726, p=0.067 2-tailed; anthropoids: F_1,12_=19.955, r^2^=0.624, p=0.001, with a considerable further improvement in fit in the latter case for a quadratic relationship, r^2^=0.760). Pooling these results using Fisher’s meta-analysis (Sokal & Rolf 1969) yields a highly significant result (χ^2^=21.99, df=4, p=0.0002), indicating that there is a consistent common trend underlying both these datasets.

**Fig. 3.**
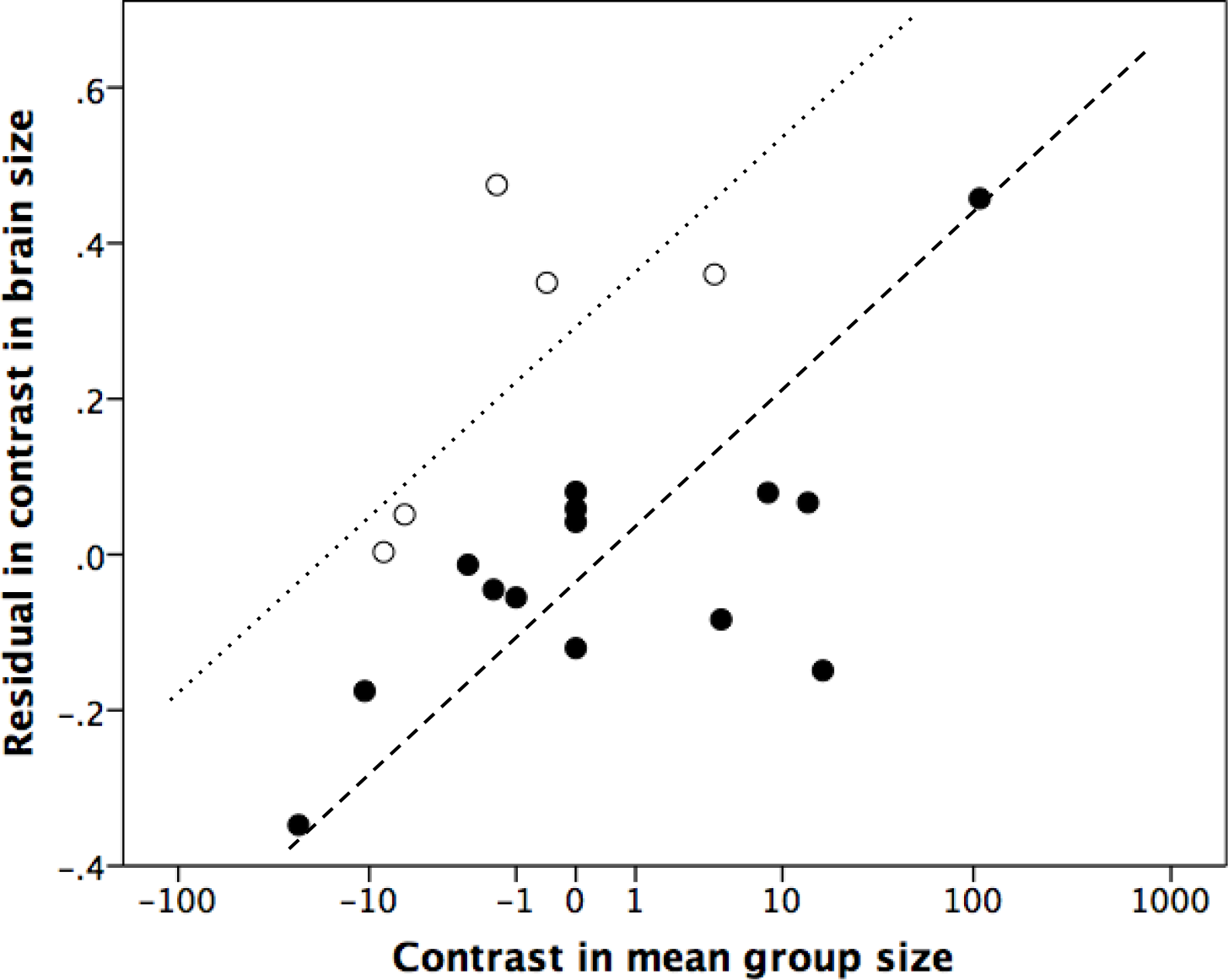
Residual of contrast in brain mass against contrast in body mass, plotted against contrasts in mean group size. Solid symbols: nodes between species pairs from the same genus; open symbols: nodes between species pairs from different genera. Hashed lines are the least squares regression lines through the two sets of data.

Given that there is an effect of group size, we perhaps need to reconsider the brain lag effect with this in mind. Fig. 4 plots the residuals from a multiple regression of contrasts in brain mass regressed on both the contrast in body mass and the contrast in group size, plotted against divergence date, for within-genus contrasts only. The multiple regression is highly significant, with significant main effects (body mass: t_16_=2.97, p=0.009 2-tailed; group size: t_16_=2.48, p=0.025 2-tailed). Since, in primates, changes in group size over phylogenetic time are almost never negative (Perez-Barberia et al. 2007), the plotted residuals are absolute values. A linear regression for these data is not significant (F_1,8_=1.70, r^2^=0.229, p=0.115 1-tailed), but a power relationship is significant (F_1,8_=4.24, r^2^=0.358, p=0.035 1-tailed, since a negative relationship is not possible). Note that the asymptote is at a residual value of ~0.035, and thus lies just above the common regression line. Mathematically, the point of inflexion is defined by the value on the X axis that is equivalent to 1/e back from the asymptotic value on the Y axis. Taking Y=0.035 as the asymptotic value and the highest datapoint as the origin (Y=0.153), this gives a value of 0.7 million years as the time it typically takes for the brain-body mass relationship to come back into balance.

**Fig. 4.**
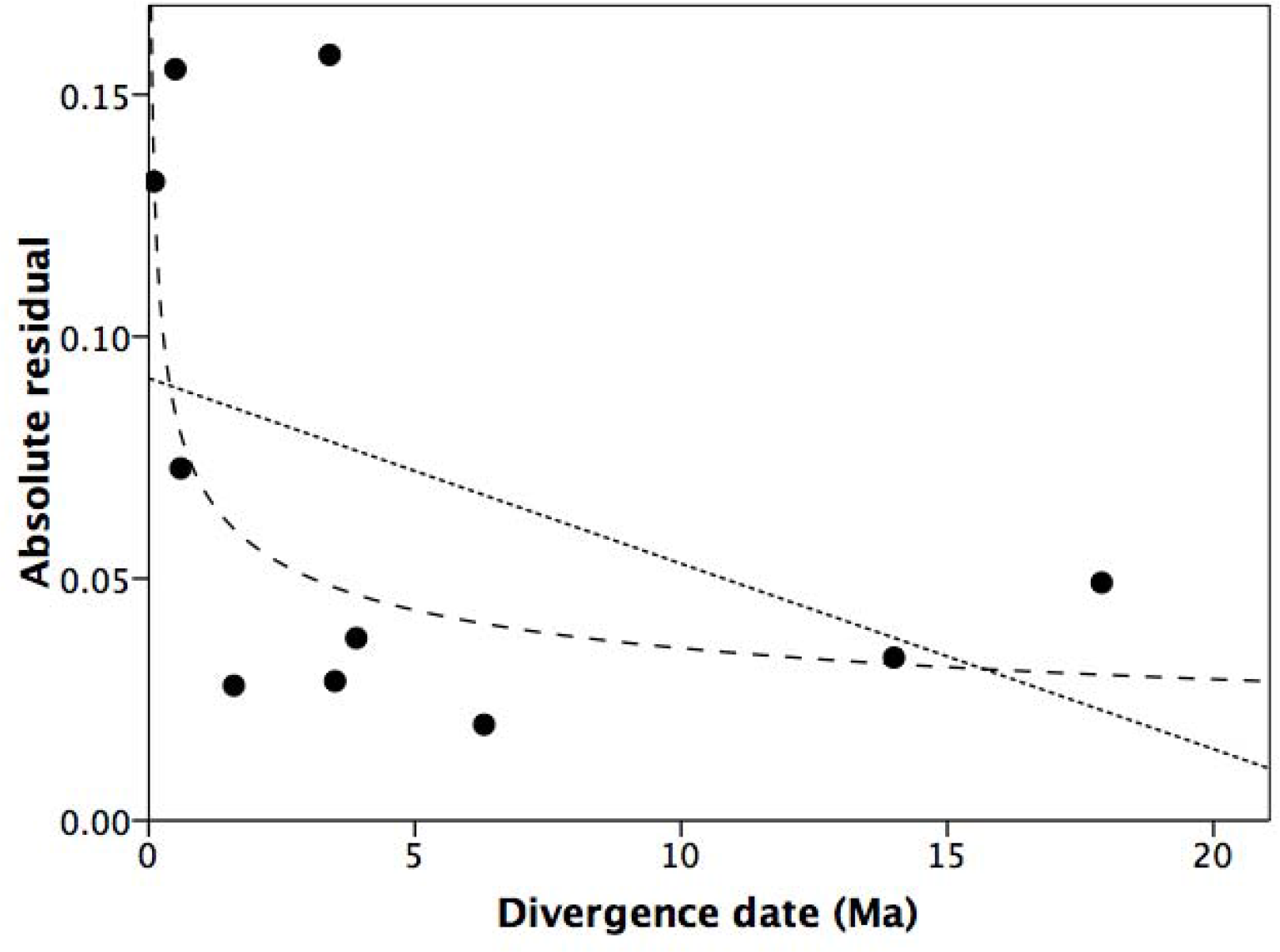
Absolute residual in contrast in brain mass against the common regression line for contrasts in both body mass and social group size, plotted against time since divergence. Linear and power regression lines are plotted.

Although the dates given by Deaner & Nunn (1999) correlate significantly (r=0.659, N=19, p=0.002) with the more recent estimates by Perelman et al. (2011), the latter tend to be deeper (the intercept for the regression equation plotting the more recent values against the older values is +2.395 Ma). Recalculation of the inflexion point for the same dataset using the Perelman et al. (2011) dates yields an estimate of 3.6 Ma. Since the Perelman et al. estimates are based exclusively on molecular data, their divergence dates identify, in effect, a last common ancestor, and thus constitute an upper limit. The date of population divergence (i.e. speciation *sensu stricto*) is likely to be a great deal less. Something in the order of 2 million years is thus probably a reasonable suggestion.

Deaner & Nunn (1997) implicitly assumed a causal relationship in which changes in body size drive changes in brain size (presumably as an inevitable consequence of the allometric relationship between the two), with changes in group size presumably being a consequence of changes in brain size (i.e. a default byproduct benefit). A path analysis of the relationship between the three variables yields a best fit model in which brain size independently predicts both body mass and group size (Fig. 5). This pattern is confirmed by a mediation analysis: body size does not significantly influence group size, or *vice versa*, via brain size. Note that by ‘predict’ here is meant ‘constrains’, and not ‘evolutionary cause’ (or driver). In evolutionary (or selection) terms, the causal arrows are reversed: increases in group size select for increases in brain size, but changes in brain size are at the same time dependent on changes in body mass to provide the sufficient energy surplus through the allometric relationship between basal metabolic rate (BMR) and body mass (Schmidt-Nielson 1984; Martin 1990) to fuel brain growth.

**Fig. 5.**
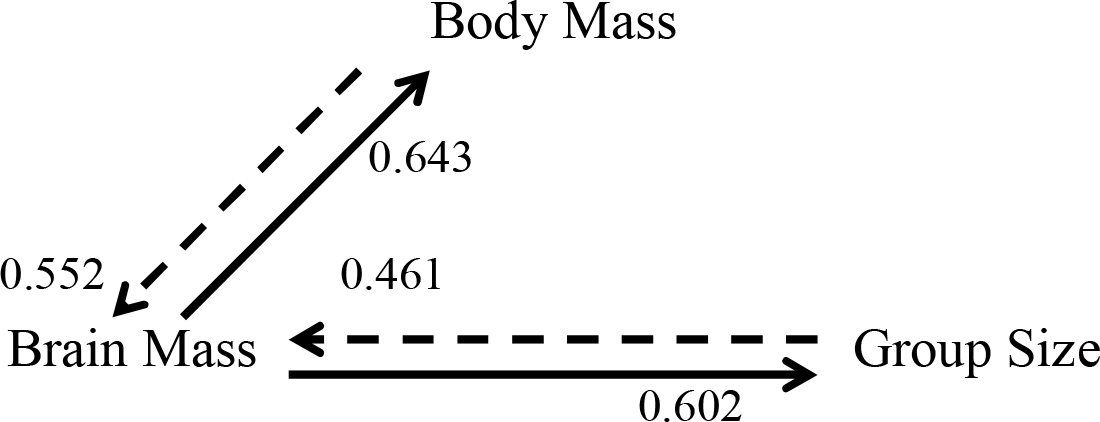
Path analysis of the functional relationships between the three main variables, contrasts in brain mass, contrasts in body mass and contrasts in mean social groups size. The plotted relationships, and the numbers given on the graph, are the significant (p<0.05) standardised slopes (ß).

Species that have undergone significant change in brain size in response to the need to increase group size will be paying an energetic cost: until body size comes back into line with brain size, they cannot benefit from the spare nutrient capacity made possible by the allometric relationship between BMR and body size (Martin 1990). To meet this demand, species will be obliged to find the additional energy and other nutrients required to fuel brain growth either through a change in diet, or by switching energy demand from other parts of the body (the expensive tissue hypothesis: Aiello & Wheeler 1995) or by reducing the energetic costs of foraging (Dunbar et al. 2009). In fact, it seems that most of the adjustment, in this sample at least, is provided by a shift to a more frugivorous (and hence more nutrient-rich) diet: on average, nodes with large residuals (absolute residual >0.05) in the contrast in brain size in Fig. 4 have a significantly greater contrast in dietary frugivory in favour of the bigger brained species than nodes with smaller residuals, although the sample is small (means of+12.7±5.8% vs -32.5±34.4%, η=0.766; F_1,5_=7.09, p=0.045 2-tailed).

## 4. Discussion

Taken together, these results imply that, as often as not, changes in group size trigger correlated changes in brain volume, initially without necessarily affecting body size (leading to high residuals); with time, body size catches up, but Fig. 4 suggests that this probably takes around 1-3 million years. The path analysis indicates that this change is invariably driven by increases in group size, and that the relationship between brain size and body size is independent of group size (i.e. is not directly determined by changes in group size). Although they endeavoured to control for ecological changes in their analysis, it seems that Deaner & Nunn (1999) failed to control for a much more important source of confound, namely the grade changes in relative brain size that occur within the primates. They also failed to control properly for social group size, mainly because they used a linear regression when the social brain relationship is explicitly non-linear (Dunbar 1992) and contains very distinct grades in the group size/brain size relationship (Dunbar 1993). Had they had more species available to them, the grade shift effects might have been lost in the error variance, but with a relatively small sample their impact is significant. Unfortunately, we are in no better position now in terms of available data than they were two decades ago because the most extensive brain dataset is still the one they used.

The initial impact of changes in brain size on the relationship between brain and body size places a significant strain on nutrient balance in species that make this change, and to balance their nutrient budget they have to increase nutrient throughput. For most monkeys, this means a more frugivorous, and less folivorous, diet. This shift to a richer diet may explain why the asymptotic value lies just above the common regression line: species that make this transition do not fully return to the common regression, but exploit their improved diet to maintain a slightly smaller body mass than would be expected. In cases, such as the *Pan-Homo* transition, where the change in brain size is massive, the increase only seems to have been possible by switching resources away from other expensive anatomical regions (notably the gut) to the brain (the expensive tissue hypothesis: Aiello & Wheeler 1995). This is, however, unlikely to be a general solution since it imposes major restrictions through gut specialisation.

This perhaps suggests a reason why several, somewhat misconceived, attempts to test the expensive tissue hypothesis on New World monkeys (Allen & Kay 2012; Hartwig et al. 2011) and mammals more generally (Navarette et al. 2011) have produced negative results: in fact, these species adopted a much simpler strategy for meeting their energy deficits, namely switching to a richer diet. In contrast, the diets of great apes are much more frugivorous than those of any monkeys (on average, 69% fruits vs 51%, N=5 and N=116 species, respectively: Campbell et al. 2007); as a result, there would have been limited room for further movement in the same direction in the transition into *Homo*, and hence a need for a more radical alternative strategy of the kind suggested by Aiello & Wheeler (1995). In fact, of course, the expensive tissue hypothesis was never offered as an explanation for primate brain evolution, but rather for brain size evolution in *Homo.* Indeed, this strategy may only be possible when the larger body mass of great apes allows more gut volume that can be spared.

In effect, it seems that Deaner & Nunn (1997) were looking at the problem the wrong way around. The assumption that the lag is based on initial changes in body size because this is physiologically and/or genetically more labile is incorrect, at least for primates. The analyses presented here suggest that quite the reverse is true: in many cases, body size change occurs *because* change has occurred in brain size, which in turn is driven by change in group size. While it is possible that the change in body size is an independent response to the same selection factor that is driving group size change (namely, predation risk: van Schaik 1982; Shultz et al. 2004; Dunbar & Shultz 2007; Bettridge & Dunbar 2012), it remains a possibility to be tested that species who opt to increase brain size ultimately need to evolve a larger body size in order to benefit from the allometric relationship between BMR and body size so as to pay for some of that increase in brain mass.

## Acknowledgments

My research is funded by an ERC Advanced Investigator grant.

